# Hybrid Derivative of Cathelicidin and Human Beta Defensin-2 Against Gram-Positive Bacteria: A Novel Approach for the Treatment of Bacterial Keratitis

**DOI:** 10.1101/2021.04.22.440925

**Authors:** Darren Shu Jeng Ting, Eunice Tze Leng Goh, Venkatesh Mayandi, Joanna M. F. Busoy, Thet Tun Aung, Mercy Halleluyah Periayah, Mario Nubile, Leonardo Mastropasqua, Dalia G. Said, Hla M. Htoon, Veluchamy Amutha Barathi, Roger W. Beuerman, Rajamani Lakshminarayanan, Imran Mohammed, Harminder S. Dua

## Abstract

Bacterial keratitis (BK) is a major cause of corneal blindness globally. This study aimed to develop a novel class of antimicrobial therapy, based on human-derived hybrid host defense peptides (HyHDPs), for treating BK. HyHDPs were rationally designed through combination of functional amino acids in parent HDPs, including LL-37 and human beta-defensin (HBD)-1 to −3. Minimal inhibitory concentrations (MICs) and time-kill kinetics assay were performed to determine the concentration- and time-dependent antimicrobial activity and cytotoxicity was evaluated against human corneal epithelial cells and erythrocytes. In vivo safety and efficacy of the most promising peptide was examined in the corneal wound healing and *Staphylococcus aureus* (ATCC SA29213) keratitis murine models, respectively. A second-generation HyHDP (CaD23), based on rational hybridization of the middle residues of LL-37 and C-terminal of HBD-2, was developed and was shown to demonstrate good efficacy against methicillin-sensitive and methicillin-resistant *S. aureus* [MIC=12.5-25.0μg/ml (5.2-10.4μM)] and *S. epidermidis* [MIC=12.5μg/ml (5.2μM)], and moderate efficacy against *P. aeruginosa* [MIC=25-50μg/ml (10.4-20.8μM)]. CaD23 (at 25μg/ml or 2x MIC) killed all the bacteria within 30 mins, which was 8 times faster than amikacin (25μg/ml or 20x MIC). After 10 consecutive passages, CaD23 did not develop any antimicrobial resistance (AMR) whereas amikacin, a commonly used treatment for BK, developed significant AMR (i.e. a 32-fold increase in MIC). Pre-clinical murine studies showed that CaD23 (0.5mg/ml) achieved a median reduction of *S. aureus* bioburden by 94% (or 1.2 log_10_ CFU/ml) while not impeding corneal epithelial wound healing. In conclusion, rational hybridization of human-derived HDPs has led to generation of a potentially efficacious and safe topical antimicrobial agent for treating Gram-positive BK, with no/minimal risk of developing AMR.

## INTRODUCTION

Infectious keratitis (IK) represents a major cause of corneal blindness worldwide, with an incidence ranging from 2.5-799 per 100,000 population-year.^1,2^ It is a common, yet potentially sight-threatening, ophthalmic emergency that often warrants hospital admission for intensive antibiotic treatment and monitoring. IK can be caused by a wide array of microorganisms, including bacteria, fungi, protozoa and viruses. Among all, bacterial keratitis (BK) has been shown to be the main cause for IK in many developed countries, including the US and the UK (>90% cases), with *Staphylococci spp.* (30-60%) and *Pseudomonas aeruginosa* (10-25%) being the two most common bacteria reported.^1,3–7^ In view of the diverse causative microorganisms, intensive broad-spectrum topical antibiotics are often administered during the initial treatment of IK. In refractory cases of IK, adjuvant therapy such as photoactivated chromophore-corneal cross-linking (PACK-CXL), amniotic membrane transplant, and therapeutic keratoplasty, are often warranted.^8–11^

The current management of IK is challenged by several factors, including the low culture yield,^1,2^ polymicrobial infection,^12,13^ and emerging trend of antimicrobial resistance (AMR).^1,14,15^ In the Antibiotic Resistance Monitoring in Ocular Microorganisms (ARMOR) 10-year prospective study, it was shown that 35-50% of the *Staphylococci spp.* were methicillin-resistant, with ~75% of them being multidrug resistant (i.e. resistant to ≥ 3 antibiotic classes).^5^ More compellingly, the increased level of AMR of the causative microorganism has been shown to negatively influence the corneal healing time and final visual outcome.^16,17^ Furthermore, complications such as corneal melt, perforation and endophthalmitis may ensue despite timely and intensive antibiotic treatment, necessitating eye-saving surgeries such as tectonic keratoplasty.^11,18,19^ All these issues highlight an unmet need for new antimicrobial treatment for IK, particularly that no new class of antibiotic has been discovered since 1987.^20^

Host defense peptides (HDPs), or previously known as antimicrobial peptides (AMPs), play vital roles in the innate immune system.^21,22^ They are a group of evolutionary conserved small molecules that are ubiquitously expressed by the immune cells and the front-line defense structures, including the ocular surface (OS).^23,24^ They have recently shown promise as potential therapeutic agents due to their broad-spectrum antimicrobial properties with minimal risk of developing AMR.^25^ These molecules are usually highly cationic and amphiphilic, with 30-50% of hydrophobicity. The cationic amino acid residues (i.e. lysine and arginine) facilitate the binding of HDPs onto the anionic bacterial membrane (*via* electrostatic interaction), while the hydrophobic residues interact with the lipid tail region of the membrane, culminating in membrane disruption, leakage of cytoplasmic contents and subsequent cell death.^26^

Previous studies have demonstrated that numerous HDPs, including LL-37, human beta-defensin (HBD)-2 and −3, are expressed at the OS and upregulated during IK.^23,27–31^ HBD-9, on the other hand, was shown to be downregulated during IK.^32^ In addition, it was demonstrated that the modulated levels of HBD-3 and HBD-9 on OS during BK returned to normal baseline following complete healing of ulcers.^33^ Furthermore, HDPs exhibit moderate *in vitro* antimicrobial activity [minimum inhibitory concentration (MIC) of around 50-100 μg/ml) against common ocular surface pathogenic isolates such as *S. aureus* and *P. aeruginosa*,^34,35^ highlighting their essential functions in human OS defense.

Despite their promising potential as effective antimicrobial therapies, several issues have impeded the successful translation of HDPs to clinical use. These include their complex structure-activity relationship (SAR), susceptibility to host / bacterial proteases, physiological conditions, and toxicity to host tissues, amongst others.^25,36,37^ In view of these issues, a number of novel strategies, including residue substitution, chemical modification, and hybridization, have been proposed to enhance the therapeutic potential of HDPs.^25,36,37^ A number of hybrid peptides, including cecropin A-melittin,^38^ cecropin A-LL37,^39^ melittin-protamine,^40,41^ amongst others,^25^ have been previously designed and reported in the literature. When compared to the parent peptide, these hybrid peptides demonstrated improved antimicrobial efficacy and/or reduced toxicity to host tissues. However, strategies in combining two human-derived HDPs from two different classes (e.g. LL-37 and HBD) have not been previously explored. In this study, we aimed to develop novel topical antimicrobial treatment for BK using hybridized HDPs derived from different combination of LL-37 and HBD-1 to −3, which are all important HDPs expressed at the OS. The in vivo efficacy and safety of the most promising molecule, CaD23 (a hybrid derivative of LL-37 and HBD-2), was further examined and validated in murine corneal wound healing and *S. aureus* keratitis models.

## RESULTS

### In vitro antimicrobial efficacy of HDPs

A total of 12 synthetic HDPs were rationally designed and synthesized based on the templates of native LL-37 and HBD-1 to −3 (**Table 1**). The MIC values (in μg/ml and μM) of all the tested antibiotics, single and hybrid peptides, in the absence and presence of 150 mM NaCl are summarized in **Table 2**.

For reference and comparison purposes, the antimicrobial efficacy of the full-length sequence of LL-37 and HBD2 and HBD3 was first determined, though extensive examination was not performed in view of the lack of translational potential for clinical use (due to the cost of synthesis associated with long length of the peptides). LL-37 demonstrated moderate-to-good efficacy against methicillin-sensitive *S. aureus* [MSSA; MIC=25.0 μg/ml], methicillin-resistant *S. aureus* [MRSA; MIC=25 μg/ml] and *P. aeruginosa* PAO1-L (MIC=50 μg/ml) whereas HBD2 and HBD3 exhibited low efficacy (MIC=100 μg/ml) against all three organisms. Subsequently, the efficacy of the three truncated versions of peptides (6-12 amino acids; based on the native template of HBD-2, HBD-3 and LL-37) and 6 first-generation hybrid HDPs (based on different combinations of LL-37, and HBD-1 to −3) were examined. All of them did not demonstrate any significant antimicrobial efficacy (MIC ≥ 200 μg/ml) against either Gram-positive or Gram-negative bacteria. Further 3 second-generation peptides (derived from CaD2 sequence) were synthesized through rational modification: (1) CaD21: substitution of cysteine with alanine; (2) CaD22: substitution of phenylalanine with tryptophan (to increase hydrophobicity); and (3) CaD23: substitution of phenylalanine and proline with tryptophan (to further enhance hydrophobicity) and substitution of cysteine with leucine. CaD21 and CaD22 demonstrated slight improvement in the antimicrobial efficacy whereas CaD23 exhibited good antimicrobial efficacy against MSSA (MIC=12.5-25.0 μg/ml), (MRSA; MIC=25 μg/ml) and methicillin-sensitive *Staphylococcus epidermidis* [MSSE; MIC=12.5 μg/ml], highlighting the importance of increased hydrophobicity for antimicrobial efficacy (**Table 1**). Moderate efficacy of CaD23 was observed against *P. aeruginosa* (MIC=25-50 μg/ml). When tested in the presence of physiological tear salt concentration (150 mM NaCl), the MIC of CaD23 against MSSA and MRSA increased by 2- to 4-fold, remained unchanged for MSSE, and remained relatively stable (1- to 2-fold increase) against *P aeruginosa*.

### In vitro cell viability and cytotoxicity

Amikacin and CaD2 did not demonstrate any lethal effect on HCE-2 cell viability at 200 μg/ml (**Figure 1A**). The IC_50_ (concentration that inhibits 50% of the cell viability) of CaD21, CaD22 and CaD23 were >200 μg/ml, >200 μg/ml and 54.6 ± 11.7 μg/ml, respectively. This demonstrated that the increased hydrophobicity with tryptophan residues enhanced the antimicrobial efficacy of CaD23 but with increased negative effect on the cell viability of HCE-2 cells. In terms of cytotoxicity, amikacin and CaD2 did not show any sign of toxicity for HCE-2 cells at 200μg/ml and CaD23 showed 30.4 ± 7.8% cytotoxicity at 200 μg/ml (**Figure 1B**). The LC_50_ (concentration that kills 50% of the cells) of CaD23 was >200μg/ml, which yielded a therapeutic index (defined by LC_50_ divided by the MIC value) of >8 for treating *S. aureus* ATCC29213. Hemolytic assay demonstrated minimal (7.1 ± 3.0%) hemolytic activity of CaD23 at 200 μg/ml (**Figure 2**). A summary of the cell viability, cytotoxicity and hemolytic results of CaD23 is provided in **Table 3**. In addition, a summary of the therapeutic index of CaD23 (defined as IC_50_, LC_50_ or HC_50_ divided by the MIC value) is provided in **Table 4**.

**Figure 1.**
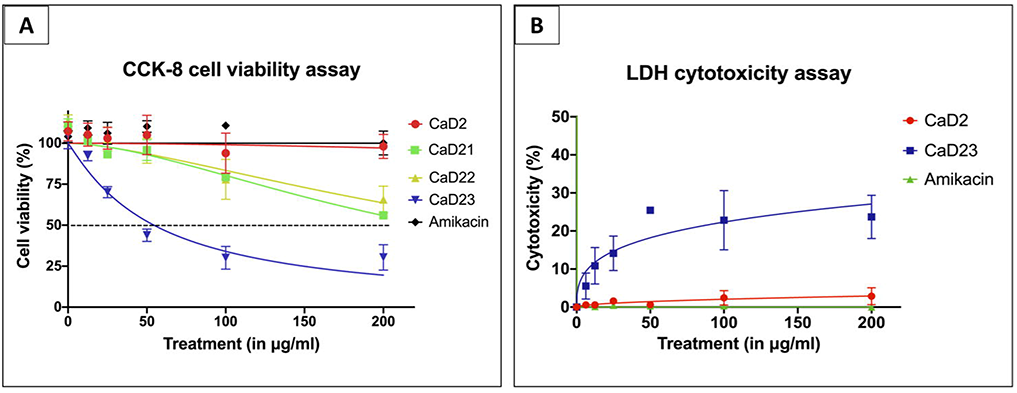
Cytotoxicity of synthetic peptides and amikacin (a commonly used antibiotic for bacterial keratitis) in various concentrations against human corneal epithelial cells (HCE-2), presented as dose-response curves (normalized, variable slope). Percentage cell viability is presented as mean ± standard deviation (depicted in error bars) of two independent experiments performed in biological duplicate. Some error bars are missing due to small standard deviation values. **(A)** Cell viability assay (using cell counting kit-8 assay) demonstrating normal metabolic activity of epithelial cells in CaD2 and amikacin but reduced activity in CaD23 (IC_50_ = 54.6 ± 11.7 μg/ml) after 3 h of treatment. IC_50_ (concentration of treatment inhibiting 50% of cell viability) is shown in a black dotted line. **(B)** Cytotoxicity assay (using lactate dehydrogenase assay) demonstrating no sign of cytotoxicity of epithelial cells in amikacin and CaD2, and low level of cytotoxicity in CaD23 (30.4 ± 7.8% at 200 μg/ml; LC_50_ > 200 μg/ml) after 3 h of treatment.

**Figure 2.**
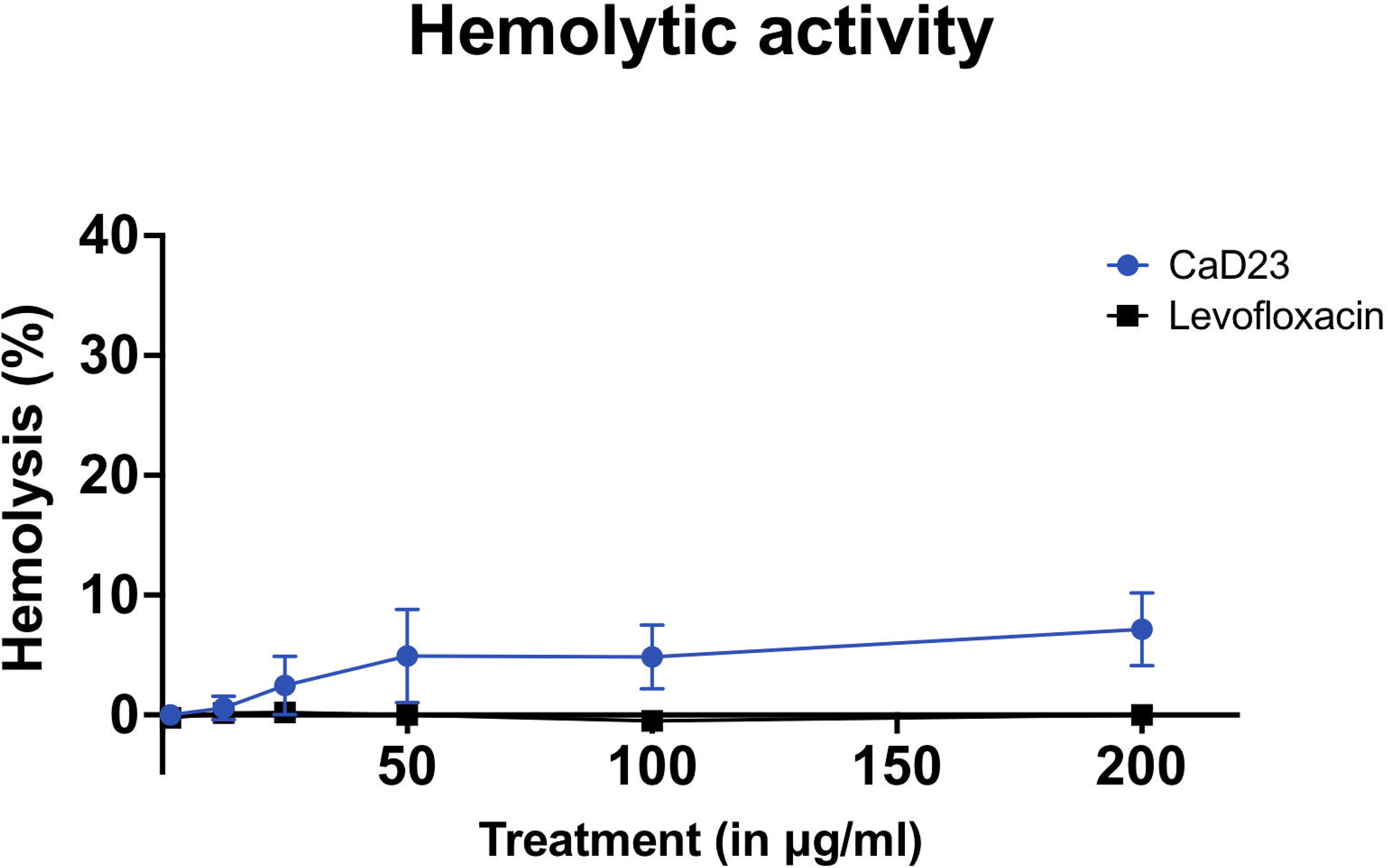
Hemolytic effect of CaD23 and levofloxacin (a commonly used antibiotic for bacterial keratitis) in various concentrations against fresh human erythrocytes, determined after 1 hour of treatment. Percentage hemolysis is presented as mean ± standard deviation (depicted in error bars) of two independent experiments performed in biological duplicate. Some error bars are missing due to small standard deviation values. The graph demonstrating minimal hemolytic effect of CaD23 against fresh human erythrocytes (only 7.1 ± 3.0% at 200 μg/ml).

### Time-kill kinetics

Time- and concentration-dependent antimicrobial activity of CaD23 was determined against MSSA (SH1000) using time-kill kinetics assay. When CaD23 was used at 2x MIC (25 μg/ml) against SH1000, it was able to achieve significant killing (99.9% or 3 log_10_ CFU/ml reduction) with 15 mins and complete killing (100%) within 30 mins of treatment (**Figure 3**). In contrast, amikacin (used at 20x MIC; 25 μg/ml) could only achieve significant and/or complete killing (99.9-100%) of SH1000 within 4 hours of treatment, which was 8 times slower than CaD23. The antimicrobial efficacy of CaD23 (25 μg/ml) and amikacin (both 10 μg/ml and 25 μg/ml) were maintained at 24 hours’ time-point, with no evidence of bacterial re-growth. Similar findings were observed when CaD23 and amikacin were tested against SH1000 in physiological tear salt concentration (150mM NaCl). CaD23 (100 μg/ml; 4x MIC) was able to achieve complete killing of *S. aureus* within 15 mins of treatment whereas amikacin (20 μg/ml; 8x MIC) was only able to achieve complete killing within 4 hours of treatment (**Figure 4**).

**Figure 3.**
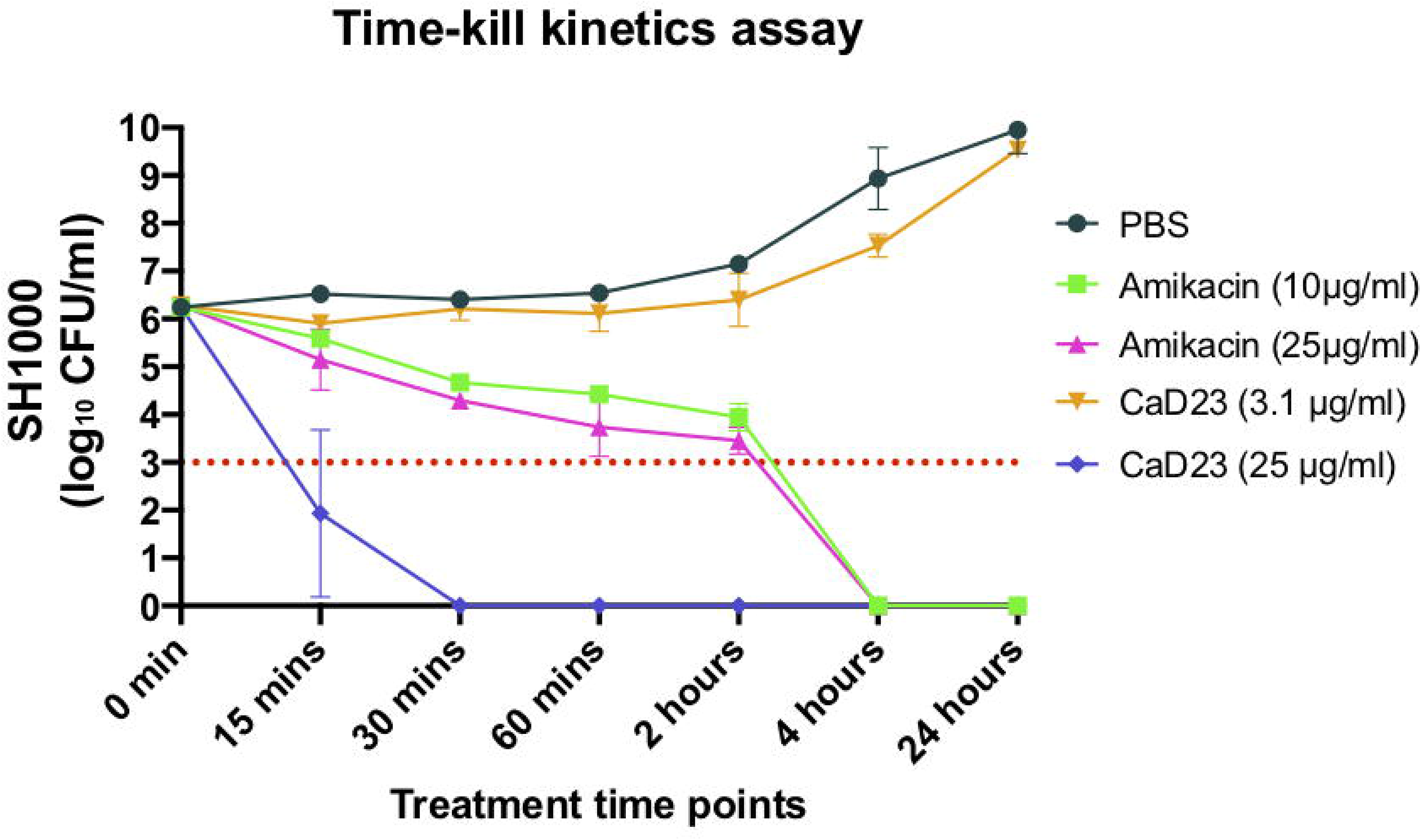
Time-kill kinetics of CaD23 (0.25x MIC and 2x MIC) amikacin (8x and 20x MIC) against *S. aureus* (SH1000) in cation-adjusted Muller-Hinton broth over 24 hours. Phosphate buffer solution (PBS) group serves as the untreated control. “0 min” represents the starting inoculum, which is around 6 log_10_ CFU/ml. The red dotted horizontal line at 3 log_10_ CFU/ml signifies the threshold of significant bacterial killing (defined as 99.9% or 3 log_10_ CFU/ml reduction of the bacterial viability compared to the starting inoculum). Data is presented as mean ± standard deviation (depicted in error bars) of two independent experiments performed in biological duplicate. CaD23 (2x MIC) was able to achieve complete (100%) killing of SH1000 within 30 mins of treatment whereas amikacin (8x MIC and 25x MIC) was only able to achieve complete killing of SH1000 within 4 hours of treatment. The antimicrobial efficacy of CaD23 and amikacin was maintained at 24 hours post-treatment.

**Figure 4.**
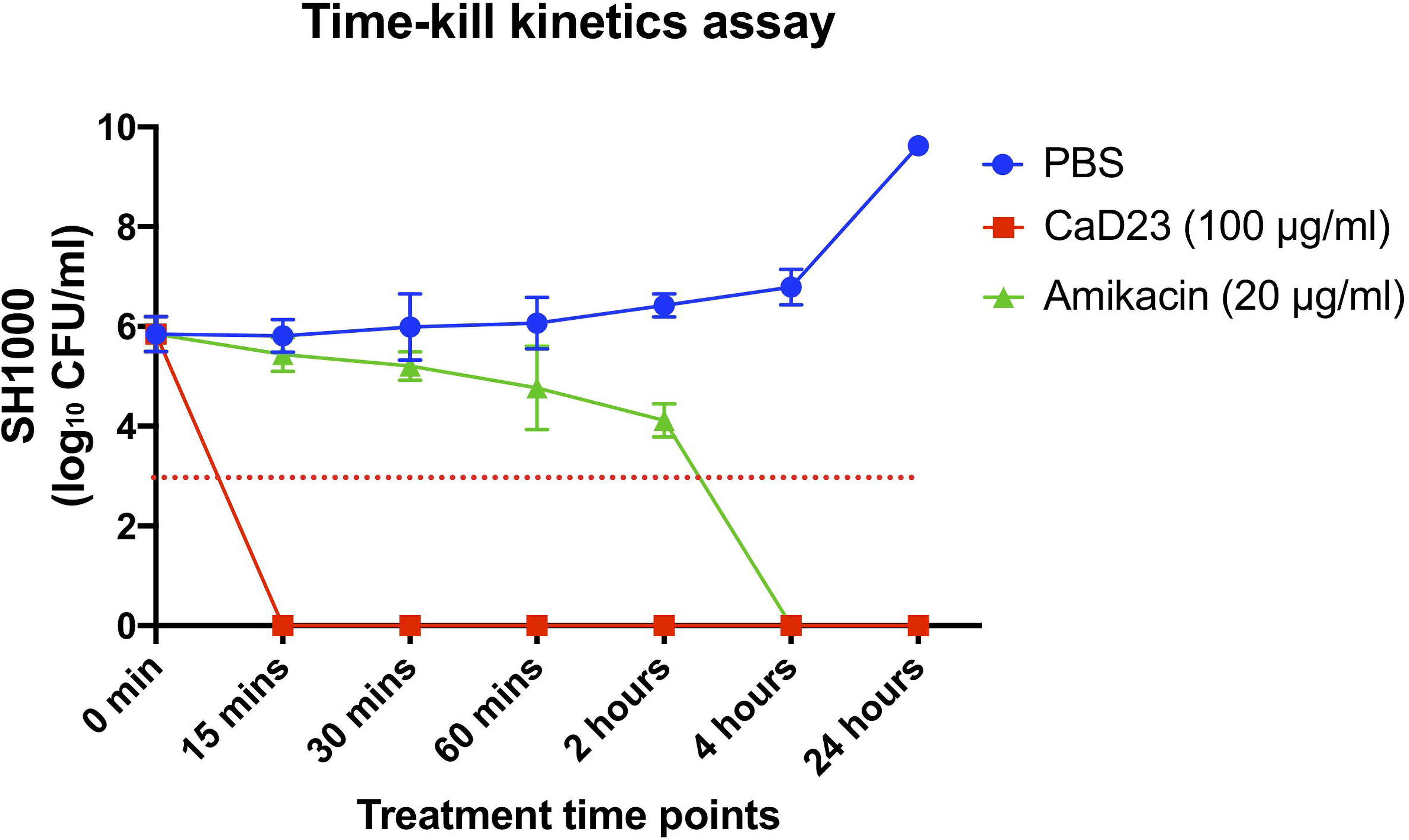
Time-kill kinetics of CaD23 (4x MIC) and amikacin (8x MIC) against *S. aureus* (SH1000) in cation-adjusted Muller-Hinton broth and physiological tear salt concentration (150mM NaCl) over 24 hours. Phosphate buffer solution (PBS) group serves as the untreated control. “0 min” represents the starting inoculum, which is around 6 log_10_ CFU/ml. The red dotted horizontal line at ~3 log_10_ CFU/ml signifies the threshold of significant bacterial killing (defined as 99.9% or 3 log_10_ CFU/ml reduction of the bacterial viability compared to the starting inoculum). Data is presented as mean ± standard deviation (depicted in error bars) of two independent experiments performed in biological duplicate. CaD23 (4x MIC) was able to achieve complete (100%) killing of SH1000 within 15 mins of treatment whereas amikacin (8x MIC) was only able to achieve complete killing of SH1000 within 4 hours of treatment. The antimicrobial efficacy of CaD23 and amikacin was maintained at 24 hours post-treatment.

### Multipassage antimicrobial resistance (AMR) study

Multipassage AMR assays were performed to evaluate the risk of development of AMR of *S. aureus* ATCC SA29213 against CaD23 and amikacin over 10 consecutive passages (days). CaD23 did not develop any AMR after 10 consecutive passages whereas amikacin developed significant AMR with a 4-fold increase in the MIC after the 2^nd^ passage and 32-fold increase in the MIC after the 10^th^ passage (**Figure 5**). This highlights the therapeutic potential of CaD23 as novel antimicrobial agent in the era of emerging AMR.

**Figure 5.**
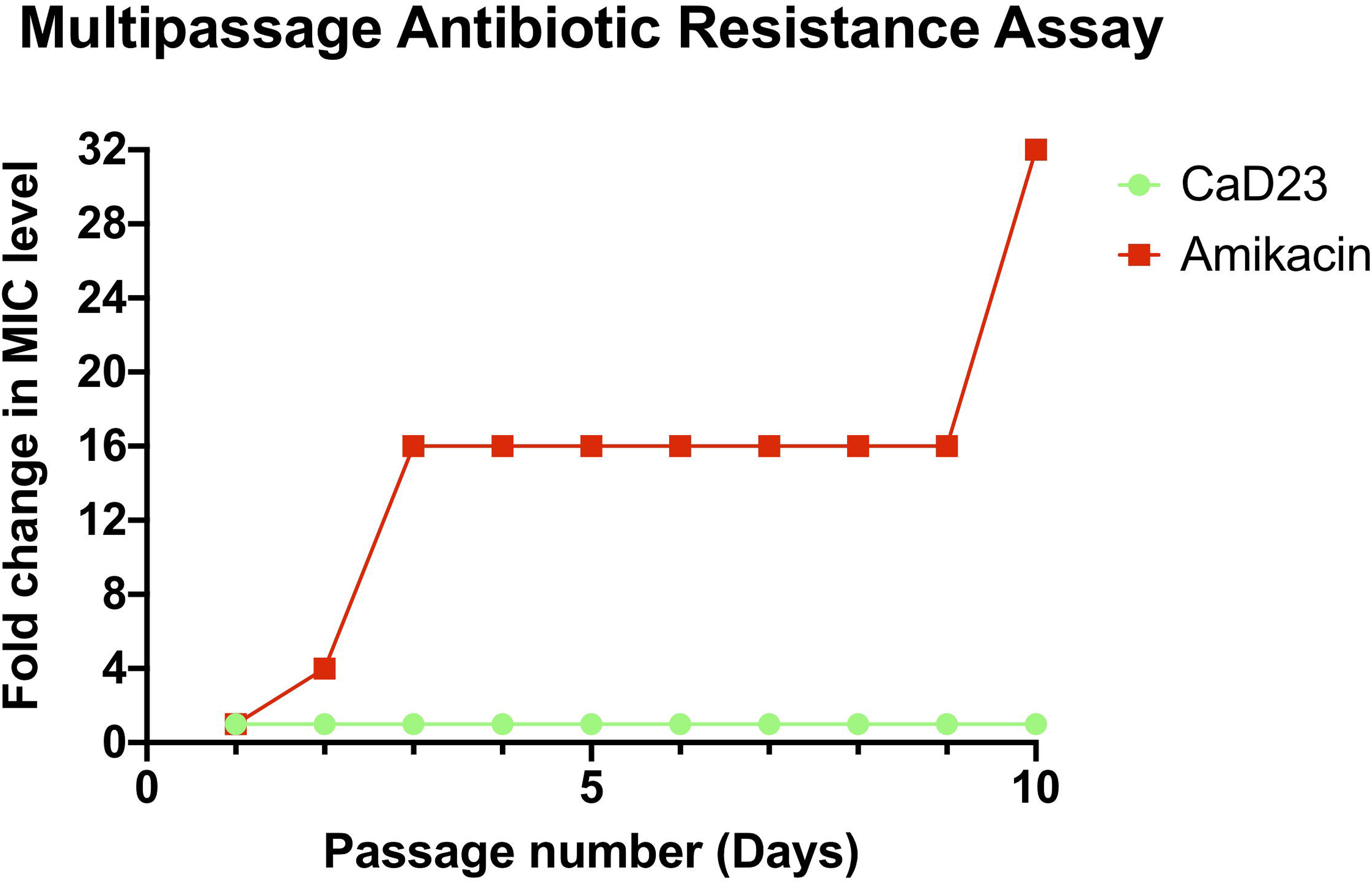
Multipassage antimicrobial resistance (AMR) assays for CaD23 and amikacin against *S. aureus* ATCC SA29213 over 10 consecutive passages (days). CaD23 did not develop any AMR after 10 passages whereas amikacin develop significant AMR with no risk of developing of AMR with a 4-fold increase in the MIC after the 2^nd^ passage and 32-fold increase in the MIC after the 10^th^ passage.

### In vivo safety of CaD23

In view of the known discrepancy between in vitro and in vivo results, the safety of CaD23 was further determined in a murine corneal epithelial wound healing model. When compared to the PBS group, both CaD23 0.03% (or 300 μg/ml) and CaD23 0.05% (or 500 μg/ml) groups did not show any significant difference in the rate of corneal re-epithelialization (all healed within 2-3 days), suggesting that both concentrations were safe for topical application (**Figure 6**). However, significant delay in corneal re-epithelialization was observed in the CaD23 0.1% group, with a mean wound size of 28.5 ± 19.9% at day 3 post-injury (p=0.004).

**Figure 6.**
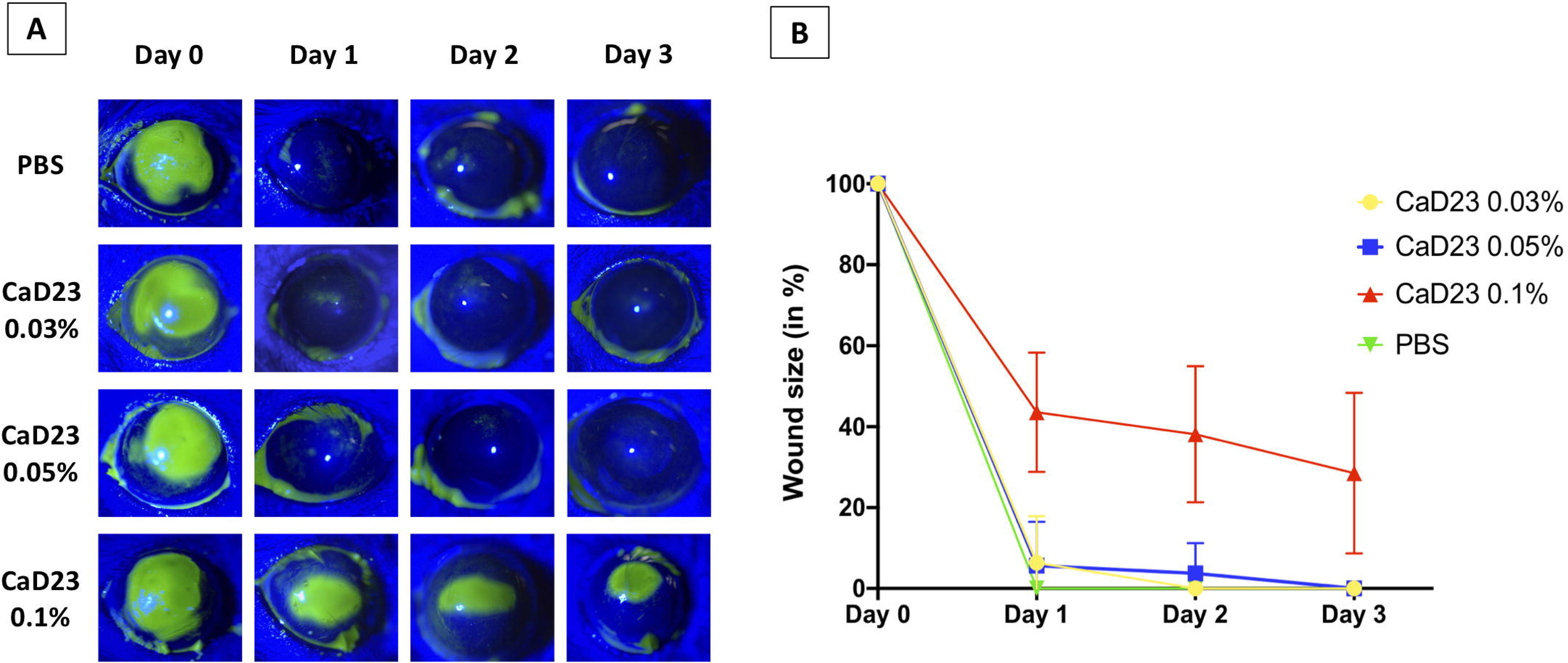
In vivo safety of CaD23 in various concentrations [0.03% (or 300 μg/ml), 0.05% (or 500 μg/ml), and 0.1% (1 mg/ml)] and phosphate buffer solution (PBS) assessed in a murine corneal epithelial wound healing model (n = 4 mice / treatment group). **(A)** Representative slit-lamp images showing the daily progress of corneal wound healing of each treatment group. The green color-stained area depicts the corneal epithelial defect. Complete corneal re-epithelialization was observed in all treatment groups, except CaD23 0.1% group, by day 3. The images were analyzed using ImageJ software (https://imagej.nih.gov/ij/).^67^ **(B)** Graphical summary of the progress of corneal re-epithelialization of each treatment group over 3 days. The corneal epithelial wound size at various time points is calculated based on the original 100% wound size at baseline. Data is presented as mean ± standard deviation.

### In vivo efficacy of CaD23

Based on the in vivo safety results, the highest tolerable concentration of CaD23 0.05% was subjected to subsequent in vivo efficacy testing in a murine *S. aureus* ATCC 29213 keratitis model. As the data was not normally distributed and was log-transformed, the results were reported in median ± interquartile range (IQR). When compared to the *S. aureus* bacterial viability in the PBS group (4.2 ± 1.3 log_10_ CFU/ml), there was a considerable reduction of bacterial viability in the CaD23 0.05% and levofloxacin 0.5% groups by 94% (3.0 ± 2.4 log_10_ CFU/ml; p=0.72) and 98% (2.4 ± 2.2 log_10_ CFU/ml; p=0.08), respectively, though statistical significance was not achieved due to wide standard deviations (**Figure 7**). At day 3 post-treatment, the clinical appearance of the corneas in the CaD23-treated group (1.2 ± 0.45; p=0.07) and the levofloxacin-treated group (1.0 ± 0.71; p=0.03) was substantially better than the untreated group (2.2 ± 0.84), based on the ocular clinical scoring (**Figure 7**).

**Figure 7.**
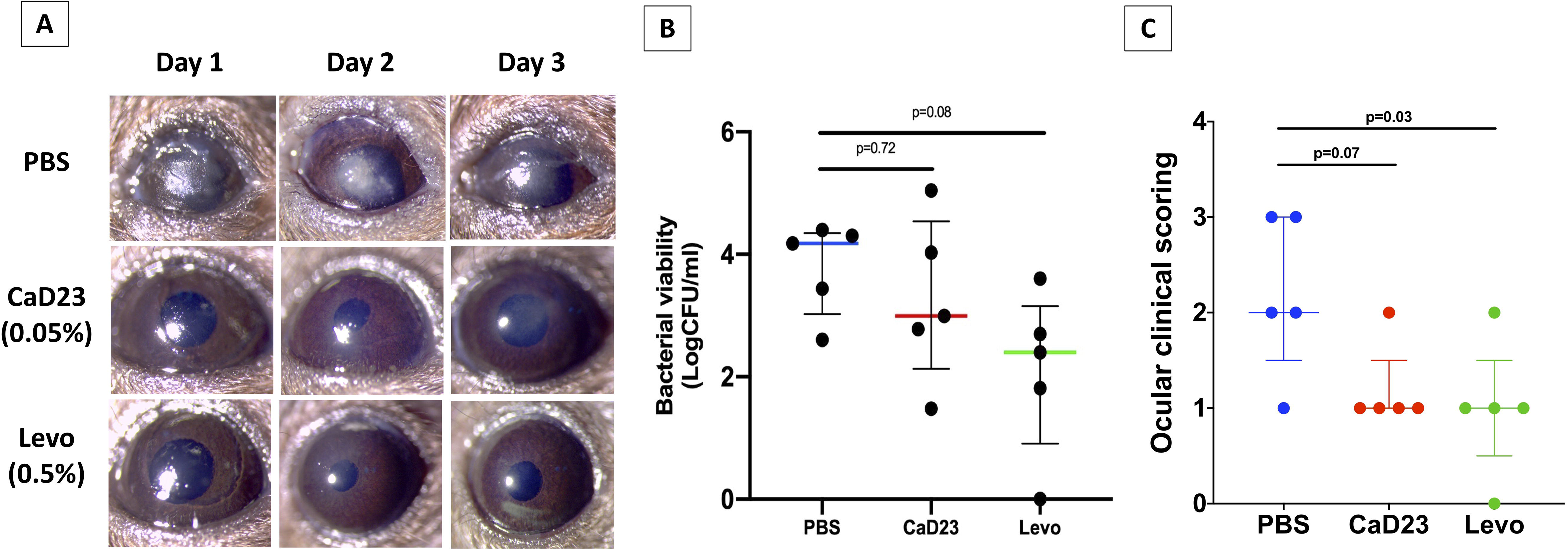
In vivo efficacy of CaD23 0.05% (500 μg/ml), levofloxacin 0.5% (positive control) and phosphate buffer solution (PBS; negative control) in a murine *S. aureus* ATCC SA29213 bacterial keratitis model (n = 5 mice / treatment group). **(A)** Representative slit-lamp images showing the corneal appearance over 3 days post-infection in each treatment group. Note the significant infiltrative changes of cornea in the PBS group as compared to the CaD23 0.05% and levofloxacin 0.5% groups. **(B)** Scatter plot showing the bacterial viability of *S. aureus* (in log_10_ CFU/ml) after 3 days of treatment. In view of the wide range of results, data is presented as median ± interquartile range. (C) Scatter plot showing the ocular clinical scoring of the clinical appearance of the *S. aureus*-infected corneas treated by different treatment. The data is presented as median ± interquartile range. The scores are interpreted as follows: 0: Clear cornea or minimal opacity, partially covering the pupil; (b) +1: Mild opacity, partially/fully covering the pupil; (c) +2: Dense opacity, partially covering the pupil; (d) +3: Dense opacity fully covering the pupil; and (e) +4: Corneal perforation or phthisis.

## DISCUSSION

IK represents a persistent and uncurbed burden on human health at a global level. Topical antibiotics are the current mainstay of treatment for BK but the efficacy is being increasingly challenged by the emergence of antimicrobial resistance.^1,2,14^ In addition, patients with BK often require long duration of treatment and sight-threatening complications may still ensue despite timely intervention,^11^ highlighting an unmet need for newer and better treatment. HDPs serve as an attractive class of antibiotics for treating IK based on the following reasons: (1) the broad-spectrum activity of HDPs can provide comprehensive coverage to a wide range of microorganisms, particularly when mixed infection is relatively common (5-20%) in IK;^1^ (2) the rapid antimicrobial action help reduce the microbial load and limit the damage to the cornea more effectively, ultimately have a better chance of preserving the vision, as well as reducing the risk of developing antimicrobial resistance, which is currently an emerging issue in ocular infection^14^; and (3) HDPs can be used as synergistic or additive agents to the current conventional antibiotics, enhancing the therapeutic index by increasing the antimicrobial efficacy and reducing the dose-related toxicity.^42,43^

In this study, we highlight a body of work in designing and developing human-derived hybrid HDP as topical treatment for BK. Hybridization strategy has been previously employed by several research groups to improve the therapeutic index of HDPs^39,40,44^ but the strategy of using human-derived hybrid HDPs (with LL-37 and HBD-2) is first of its kind. The initial concept of developing synthetic HDPs derived from LL-37 and HBD was founded on the observation of the upregulation of these key HDPs at the ocular surface during IK.^23,27,33,45^ Furthermore, LL-37 and HBD-2 and −3 were shown to exhibit good antimicrobial efficacy against a range of organisms.^34,45^ We have selected the mid-region of LL-37, consisting of residues 18-29 of LL-37 (i.e. KR-12) as part of the hybrid template as studies have demonstrated that KR-12, though much shorter than parent LL-37, exhibited similar antimicrobial activity against *E. coli* (MIC = ~64 μg/ml) with reduced toxicity to host cells.^46^ C-terminal of HBD1-3 was used as the other part of the hybrid template in view of its rich content of cationic residues, which have been shown to play an important role in interacting with anionic bacterial membrane and killing of bacteria.^47,48^

Our initial attempt of engineering single linear HDPs (including LL-37 and HBD-2 and −3) and hybrid HDPs (based on LL-37 and HBD-1 to −3) did not yield any compound with good antimicrobial efficacy. However, with systematic SAR analysis and modification, particularly through substitution of proline and phenylalanine with tryptophan residues (to increase hydrophobicity and membrane partitioning), ensued the development of peptide with enhanced antimicrobial efficacy against MSSA (MIC=12.5-25 μg/ml) and MRSA (MIC=25 μg/ml), serving as a proof-of-concept of this novel design strategy. Interestingly, increased hydrophobicity augmented the antimicrobial activity of HDP against Gram-positive bacteria more than Gram-negative bacteria. This is likely attributed to the different compositions in the bacterial membrane between Gram-positive bacteria, which consist of a thick peptidoglycan layer, and Gram-negative bacteria, which comprises an additional outer membrane with abundance of negatively charged lipopolysaccharide for which cationicity of the peptide plays a more important role.^49^ In addition, the efficacy of CaD23 (18 amino acids in length) is at least equal to or stronger than the full-length parent LL37 (MIC = 25-50 μg/ml), HBD-2 (MIC=100 μg/ml) and HBD-3 peptides (MIC = 50-100 μg/ml). Peptides with shorter sequence not only have the advantage of lower manufacturing cost but may also have a lower risk of inducing immunogenicity.^25,50^ Moreover, the antimicrobial efficacy of CaD23 is comparable to some of the HDPs that are developed for ocular surface infection, including esculentin-1a(1-21)NH2,^51^ RP444,^52^ melimine and its derivatives,^40,53^ and ɛ-lysylated Mel-4.^54^

In the time-kill kinetics study, we demonstrated that CaD23 at 2x MIC was able to achieve complete killing of *S. aureus* within 30 mins as compared to 4 hours with amikacin at 20x MIC (i.e. 8 times slower than CaD23). We also observed that increasing the concentration of amikacin from 8x MIC to 20x MIC did not expedite its anti-bacterial action against *S. aureus*, suggesting that the speed of bacterial killing is more related to the underlying mechanism of action than the concentration of antibiotic. The rapid killing action of CaD23 is likely related to a membrane perturbation effect, which is in contrast to amikacin where it exerts its anti-bacterial activity via intracellular inhibition of the 30S subunit of bacterial ribosome.^55^ Theoretically, the rapid and membrane disruptive action of CaD23 should result in a lower risk of developing antimicrobial resistance as the bacteria has less time to adapt and require substantial modification of genome to develop effective resistant mechanisms, which are shown in conventional antibiotics.^54^ Further studies examining the underlying mechanism of action of CaD23 and its tendency to develop antimicrobial resistance would be valuable.

Based on the LDH cytotoxicity assay, CaD23 was shown to be relatively non-cytotoxic (20-30% toxicity) at 200 μg/ml, with a therapeutic index of >8-16 against *S. aureus* (defined by LC_50_ concentration divided by MIC value). In addition, the hemolytic activity of CaD23 at 200 μg/ml was only less than 10%. Interestingly, the IC_50_ value (based on cell viability assay) was around 50 μg/ml, considerably lower than the LC_50_ value. This discrepancy may be due to the fact that some of the cells were metabolically inhibited but the cell membrane was not disrupted, therefore the IC_50_ value was lower than the LC_50_ value. That said, our in vivo corneal epithelial wound healing study showed that CaD23 did not demonstrate any significant toxicity when used at 500 μg/ml (or 0.05%), which was 10 times higher than the IC_50_ value. The observed discrepancy between in vitro and in vivo findings is likely attributed to the inherent dynamic environment of ocular surface with eye blinking, high tears turnover, and drainage, and dilution of the peptide by the tears.^56^ This also suggests that in vitro plate-based static assays such as cytotoxicity or cell viability assays could potentially overestimate the in vivo cytotoxicity of drugs that are developed for ocular topical application. The shortcoming of in vitro assays may be addressed by the recently developed novel ex vivo biomimetic model where it could simulate the dynamic and complex interface between the ocular surface and the external environment, allowing a better prediction of the in vivo effect.^57^

Our in vivo BK study demonstrated that CaD23 0.05% (20x MIC) was able to reduce the bacterial viability of *S. aureus* by 94% when compared to the untreated control group, which was more than the pre-defined endpoint of the study (i.e. 1 logCFU or 90% reduction in the bacterial bioburden). However, the effect was not statistically significant due to the considerably wide standard deviation observed in both the treatment and the control groups. This similarly explained the insignificant improvement in the levofloxacin-treated group, though there was 1.8 logCFU median reduction in the bacterial bioburden compared to the PBS group. Nevertheless, these results serve as a strong proof-of-concept that CaD23 may be employed as a potentially efficacious treatment for treating Gram-positive bacterial keratitis, but a larger sample size will be required to fully ascertain its efficacy. In addition, we had chosen to sacrifice the mice at day-3 post-infection because *S. aureus* keratitis had been shown to be most severe in C57BL/6J mice at 3-day post-infection, as compared to other stains of mice such as BALB/c and A/J mice. In addition, *S. aureus* keratitis has been shown to improve spontaneously in C57BL/6J mice at day-5 post-infection and beyond.^58^ To investigate the longer-term in vivo antimicrobial efficacy of CaD23 in *S. aureus* keratitis, other strains of mice may be required to examine this aspect.

One potential approach to improve the therapeutic effect of HDP-derived antimicrobial treatment is to use them in combination with antibiotics as peptide-antibiotic synergism has been demonstrated in several studies.^42,43,59^ This attractive antimicrobial strategy not only helps extend the lifespan and broaden the antibacterial spectrum of conventional antibiotics, but also reduce the dose-dependent toxicity associated with HDPs and antibiotics.^60^ Recently, our group has also shown that FK16, a cathelicidin-derived molecule, could improve the antimicrobial efficacy of vancomycin, a glycopeptide antibiotic, against PA by 8-fold.^42^ This is likely ascribed to the different mechanisms of action between FK16 and vancomycin where the membrane disruptive action of FK16 facilitates the diffusion of vancomycin across the bacterial membrane. Hence, future work investigating the potential synergism between CaD23 and commonly used antibiotics for BK would be useful.

At present, our aim is to develop the human-derived hybrid HDPs as topical treatment for BK. However, it is interesting to note that the hemolytic activity of CaD23 was very low (7.1% at 200 μg/ml), suggesting that it can potentially be developed for treating systemic infection. It also reduces the concern of systemic toxicity when it is applied to the ocular surface because systemic absorption of ocular topical treatment can occur, particularly when the drugs are hydrophobic.^56^ The considerable disparity between the hemolytic activity and cytotoxicity against corneal epithelial cells could be attributed to the difference in the membrane structure (particularly the cytoskeleton) and the metabolic structures of the non-nucleated cells (e.g. red blood cells) and nucleated cells (e.g. corneal epithelial cells).^61^ Further studies examining the efficacy and stability of CaD23 in serum as well as its interaction with blood proteins will be conducted to determine its therapeutic potential for treating systemic infection.

In conclusion, we demonstrated that rational hybridization of LL-37 and HBD-2 serves as a useful strategy in translating the therapeutic potential of human-derived HDPs. Future work examining the efficacy and safety of combined CaD23-antibiotic therapy would be beneficial. Potential strategies such as N- and C-terminal modifications, introduction of unnatural amino acids and nanoformulation of CaD23 will be further explored with an aim to enhance the antimicrobial efficacy, reduce the toxicity and improve the stability of the peptide.^25^

## MATERIALS AND METHODS

The commercially synthesized peptides and commonly used antibiotics for IK were first examined for their in vitro antimicrobial efficacy against a range of bacteria. In vitro cytotoxicity of these antimicrobial agents was then determined against human corneal epithelial cells (HCE-2, CRL-11135, ATCC, UK) using cell viability assay and cytotoxicity assay, and against human erythrocytes using hemolytic assay. The most promising synthetic peptide, CaD23, was further examined for its time- and concentration-dependent in vitro antimicrobial activity. Finally, in vivo efficacy and safety of CaD23 were evaluated in corneal wound healing and *S. aureus* keratitis murine models. All the assays described in this study were conducted in biological duplicate and in at least two independent experiments, with appropriate positive controls (PCs) and negative controls (NCs). Continuous values were expressed in mean ± standard deviation (SD), unless specified otherwise.

### Design and synthesis of HDPs

A template-based design method was used to design our human-derived HDPs. The native peptide sequences were obtained from an established protein bank database (https://www.uniprot.org/). Several human-derived HDPs, specifically HBD-1, −2, and −3, and cathelicidin (LL-37), were subject to testing and hybridization. The physicochemical properties of the designed peptides, including peptide weight, net charge, hydrophobicity (<H>), and amphiphilicity / hydrophobic moment (< μH>, were analyzed using a computational programmes such as PepCalc (https://pepcalc.com/) and HeliQuest (http://heliquest.ipmc.cnrs.fr). Various strategies, including residue substitution and hybridization, were employed for SAR analysis and for improving the therapeutic index (i.e. increasing the antimicrobial efficacy and reducing the toxicity profile). The native and synthetic peptide sequences are shown in **Table 1**.

Synthesis of the single and hybrid HDPs was based on the knowledge of the functional regions of the native templates. Three single, short-sequenced (truncated) peptides, based on the native template of HBD2, HBD3 and LL-37, were first generated. Truncated versions of the parent peptides were engineered as this strategy has been shown to serve as a useful method in improving the efficacy of peptides and reducing the cost of synthesis, which represents a significant translational barrier of peptide-based antimicrobial therapy.^25^ The C-terminal region of HBD2 and HBD3 was synthesised and examined in view of the presence of high cationicity (i.e. rich of lysine and/or arginine residues), which are important for the antimicrobial efficacy.^26,47,48^ In addition, the middle region of LL-37, same as the KR12 molecule, was synthesized as it has been shown to exhibit efficacy equivalent to the full-length of LL-37.^46^

All antibiotics, including amikacin (an aminoglycoside) and levofloxacin (a fluoroquinolone), were purchased from Sigma-Aldrich, United Kingdom. Both antibiotics were used as positive controls as they were commonly used for treating BK.^1,4,62^ The full-length peptides were commercially produced by Anaspec (Cambridge, UK; for LL-37) and PeproTech (London, UK; for HBD-2 and HBD-3). All other peptides were commercially produced by Mimotopes (Mulgrave Victoria, Australia) via the traditional solid phase Fmoc synthesis method. All the synthetic peptides were purified by reverse-phase high performance liquid chromatography (RP-HPLC) to >95% purity and characterized by mass spectrometry.

### Range of microorganisms being tested

A range of Gram-positive and Gram-negative laboratory- and clinical-strain bacteria were used for the experiments. These included laboratory-strain methicillin-sensitive *S. aureus* (MSSA; including SH1000 and ATCC SA29213), laboratory-strain MRSA (ATCC MRSA43300), clinical-strain MRSA (MRSA-OS; an IK isolate), laboratory-strain methicillin-sensitive *Staphylococcus epidermidis* (MSSE; ATCC SE12228), laboratory invasive-strain *P. aeruginosa* (PAO1-L), and clinical cytotoxic-strain *P. aeruginosa* (PA-OS; an IK isolate).

### Determination of antimicrobial efficacy using MIC assay

In vitro antimicrobial efficacy of the antibiotics and designed HDPs was determined using an established MIC assay with broth microdilution method approved by the Clinical and Laboratory Standards Institute (CLSI).^63^ Briefly, the microorganisms were cultured on Tryptone Soya Agar (TSA) and incubated overnight for 18-21 hours at 37 °C. Bacterial inoculums were subsequently prepared using the direct colony suspension method.^63^ Three to five bacterial colonies were obtained from the agar plate and inoculated into an Eppendorf tube containing 1 ml of cation-adjusted Muller-Hinton broth (caMHB), consisting of 20-25 mg/L calcium ions (Ca^2+^) and 10-12.5 mg/L magnesium ions (Mg^2+^). The bacterial suspension was adjusted to achieve a turbidity equivalent to 0.1 OD_600_ or 0.5 MacFarland, containing ~1.5 × 10^8^ colony-forming unit (CFU)/ml, which was then further diluted in 1:150 in caMHB to reach a final bacterial concentration of ~1×10^6^ colony forming units (CFU)/ml. Each treatment (peptide or antibiotic) was prepared in 1:2 serial dilution in 96-well polypropylene microplates (with a final treatment volume of 50 μl per well), followed by the addition of 50 μl of 1×10^6^ CFU/ml bacteria into each well (with a final bacterial concentration of 5×10^5^ CFU/ml). As the HDPs are known to be influenced by the salt content,^25^ the MIC assay was also performed in the presence of physiological tear salt concentration (150 mM NaCl). The MIC values, defined as the lowest concentration of the antimicrobial agent that prevented any visible growth of bacteria, were determined after 24 hours of incubation with treatment.

### Cell viability and cytotoxicity assays

Cell viability and cytotoxicity of antibiotics and peptides were determined against human corneal epithelial cells (HCE-2, CRL-11135, ATCC, Manassas, Virginia, USA) using cell-counting-kit-8 (CCK-8) assay (Sigma Aldrich, Merck Life Science UK Limited, Dorset, UK) and lactate dehydrogenase (LDH) assay (ThermoFisher Scientific, UK), respectively, as per manufacturer’s guidelines. HCE-2 cells were cultured and seeded into a 96-well plate at a density of 7.5 × 10^3^ cells/well and allowed to attach overnight and grew to 80-90% confluency in keratinocyte serum free medium (KSFM) supplemented with human recombinant epidermal growth factor, bovine pituitary extract, hydrocortisone, and insulin. Once the confluency reached 80-90%, the cells were incubated with treatment for 3 hours before OD_450_ measurement was taken using BMG Clariostar microplate reader (BMG Labtech Ltd., Aylesbury, UK). Appropriate controls were used, including 0.1% Triton X-100 as positive control and KSFM as negative control. Cell viability was calculated using the following formula: [((I_treatment_ – I_NC_) / (I_NC_)) × 100; I=intensity] and cytotoxicity was calculated using the following formula: [((I_treatment_ – I_NC_) / (I_PC_ – I_NC_)) × 100].

### Hemolytic assay

Ethical approval was obtained from the Local Research Ethics Committee of University of Nottingham prior to the experiment (Ref: 176-1812). The experiment was conducted in accordance with the relevant guidelines and regulations. Hemolytic assay was performed according to the previously established protocol.^42^ Briefly, 5 ml of fresh human blood was collected from healthy participants with written informed consent, in an EDTA tube and centrifuged for 10 mins at 1300 g at 10 °C for separation of plasma. The remaining erythrocytes were rinsed and centrifuged further three times in Ca^2+^/Mg^2+^ free Dulbecco’s phosphate-buffered saline (DPBS). Subsequently, erythrocytes were diluted to 8% v/v in DPBS and incubated with 100 μl of treatment, positive control (1% Triton X-100) and negative control (DPBS), all in 1:1 ratio, for 1 hour (final erythrocytes concentration = 4% v/v). After 1 hour of incubation, the plate was centrifuged at 500 × g for 5 mins and 100 μl of supernatant of each well was transferred into a 96-well plate for measurement at OD_540_. Hemolysis (%) was calculated as [(I_treatment_ – I_NC_) / (I_PC_ – I_NC_)] × 100.

### Time-kill kinetics assay

Time-kill kinetics assay was performed to determine the time-dependent and concentration-dependent in vitro antimicrobial effects of the CaD23 and amikacin, a commonly used topical antibiotics for BK, against 100 μl of ~1 × 10^6^ CFU/ml of SH1000 (1:1 treatment-bacteria ratio) at various time points, including 0 min (pre-treatment), 15 mins, 30 mins, 60 mins, 2 hours, 4 hours, and 24 hours. At each time point, 10 μl of the treated bacteria was transferred to an Eppendorf tube containing 90 μl of PBS, which was then serially diluted in 1:10 concentration for inoculation on agar plates and incubated overnight for 18-21 hours at 37 °C for enumeration of CFU.

### Multipassage antimicrobial resistance (AMR) study

Multipassage AMR assays were performed to evaluate the risk of development of AMR of *S. aureus* (ATCC SA29213) against CaD23 and amikacin over 10 consecutive passages (days). The assay was conducted in a similar manner of the MIC assay described above. After determining the initial MIC level of each treatment at baseline (passage 1), bacterial suspensions were obtained from the 0.5x MIC well of each treatment and were adjusted in caMHB to achieve a turbidity equivalent to 0.1 OD_600_ (containing ~1.5 × 10^8^ CFU/ml). This was then further diluted in 1:150 in caMHB to reach a final bacterial concentration of ~1×10^6^ colony forming units (CFU)/ml. Subsequently, 50 μl of 1×10^6^ CFU/ml of the bacteria was added to the corresponding serially diluted treatment and the MIC level was determined after 24 hours of incubation with treatment. Development of AMR was defined as ≥4-fold increase in the MIC level compared to the baseline.

### In vivo efficacy and safety studies

The in vivo studies were conducted in two stages, namely the corneal wound healing study (for safety) and BK study (for efficacy), based on previously established protocols.^54,64^ Wild-type C57BL/6J mice (8-9 weeks old, male, average weight of 25g) were used in view of the consistent and reproducible results demonstrated in previous studies.^54,64^ The mice were maintained and treated in compliance with the Guide for the Care and Use of Laboratory Animals (National Research Council) and the ARVO statement for the Use of Animals in Ophthalmic and Vision Research. All animal studies were conducted at the Singapore Eye Research Institute, Singapore, and were approved by the Animal Welfare & Ethical Review Body (AWERB), University of Nottingham, UK (Ref: UoN-Non-UK #16), the Institutional Animal Care and Use Committee (IACUC) and the Institutional Biosafety Committee (IBC) of SingHealth, Singapore (Ref: 2019/SHS/1491). All experiments were performed in accordance with relevant guidelines and regulations. General anesthesia was administered using intraperitoneal injections of xylazine (10 mg/kg) and ketamine (80 mg/kg)], and a drop of topical proxymetacaine hydrochloride 0.5% was administered immediately before wounding and/or infecting the corneas. Upon completion of the experiments, all mice were sacrificed according to the method of humane killing set out in Animals (Scientific Procedures) Act 1986 Schedule 1 using overdose of general anesthesia via intraperitoneal route. The study was reported in accordance with ARRIVE guidelines (https://arriveguidelines.org).^65^

#### In vivo corneal wound healing study

Drug drainage, blinking and tear film were considered during dosing translation from in vitro to in vivo use.^52^ Based on the MIC of CaD23 against *S. aureus* ATCC 29213 (= 25 μg/ml), a range of concentration of CaD23 was chosen, including 300 μg/ml (0.03%; 12x MIC), 500 μg/ml (0.05%; 20x MIC) and 1mg/ml (0.1%; 40x MIC).

In vivo safety of CaD23 (in 0.03%, 0.05% or 0.1%), and PBS (negative control) was first determined in a mouse corneal epithelial wound healing model. In the absence of *in vivo* pilot data of our designed HDPs, the sample size was calculated based on a previous study.^66^ This was designed as a non-inferiority trial to ensure that the HDPs did not affect the wound healing when compared to PBS (control). The non-inferiority margin was set at 10% difference of the wound size between HDPs and PBS at 3 days (deemed as significantly different), with a standard deviation (SD) of 5% (Cohen’s d=2.0), power=80% and p<0.05. A minimum sample size of 4 mice / treatment group was needed.

All mice were randomly allocated to each of the four treatment groups (n=4 mice/group). Prior to the study, all eyes were examined with slit-lamp biomicroscopy to confirm the health of corneas. Under general and topical anesthesia, the central 2 mm corneal epithelium was gently debrided with sterile Beaver mini-blades, leaving the basal lamina intact. Each treatment was applied immediately after wounding, then 4 times a day at 3-hour interval for 3 days (total dose of treatment per mouse = 14). Corneal epithelial defect was assessed using a cobalt-blue filter-equipped slit-lamp biomicroscopy and photography with staining with topical sodium fluorescein 1% at baseline (immediately post-debridement) and daily up to 3 days post-treatment. Fluorescein-stained images of the corneal wound defect were analyzed using ImageJ software (https://imagej.nih.gov/ij/).^67^ The main outcome measure was the wound size at the end of day 1, 2 and 3 [expressed as % of the original wound size in mean and standard deviation (SD)]. Difference in the wound size between groups was analyzed using one-way ANOVA with Dunnett’s post hoc test (PBS as the control group).

#### In vivo S. aureus keratitis study

Based on the in vivo safety data, the highest tolerable concentration of CaD23, 500 μg/ml (0.05%), was used in the subsequent *S. aureus* keratitis murine model. Levofloxacin 0.5%, a commonly used antibiotic for BK in clinical setting,^4^ and PBS were used as the positive and negative controls, respectively. In the absence of *in vivo* pilot data, the sample size was calculated based on a previous study.^64^ To detect an effect size of 1 LogCFU (or 10 times) difference in the bacterial load (significant antimicrobial efficacy) between HDPs (mean=5 logCFU) and PBS (mean=6 logCFU; NC), with a SD of 0.5 logCFU (Cohen’s d=2.0), power=80% and p<0.05, a minimum sample size of 4-5 mice/group is required.

All mice were randomly allocated to each treatment group (n=5 mice/group). Slit-lamp examination was performed before the start of experiment to confirm the health of corneas. Under general and topical anesthesia, the central 2 mm corneal epithelium was gently removed with sterile Beaver mini-blades. 10 μl of ~1×10^8^ CFU/ml of ATCC SA29213 was applied topically onto the cornea and the lid was held shut for 1 min. At 6 hours post-infection, 10 μl of treatment was applied directly onto the infected corneas with a dose regimen of 4 times a day at 3-hour interval for 3 days (total dose of treatment per mouse = 12). The eyes were monitored daily using slit-lamp biomicroscopy and photography. Ocular clinical scoring was adapted from a previous method described by Clemens et al.,^52^ with minor modifications: (a) 0: Clear cornea or minimal opacity, partially covering the pupil; (b) +1: Mild opacity, partially/fully covering the pupil; (c) +2: Dense opacity, partially covering the pupil; (d) +3: Dense opacity fully covering the pupil; and (e) +4: Corneal perforation or phthisis.

At the end of day 3, all animals were sacrificed, and the infected eyes were enucleated. The whole corneas were subsequently dissected and homogenized in 1 ml of sterile PBS using sterile glass micro-beads. The homogenised infected corneal tissue suspension was serially diluted in 1:10 and plated on TSA plates in triplicates for enumeration of CFU after 24 hours incubation at 37 °C. The main outcome measures were the ocular clinical scoring and the residual bacterial load at 3-day post-treatment (expressed as log_10_ CFU/ml, which was the same as log_10_ CFU/cornea) and the difference among groups was analyzed using one-way ANOVA with Dunnett’s post hoc test (PBS as the control group).

## References

1. Ting DSJ, Ho CS, Deshmukh R, Said DG, Dua HS. Infectious keratitis: an update on epidemiology, causative microorganisms, risk factors, and antimicrobial resistance. Eye (Lond). 2021;35(4):1084–101.

2. Ung L, Bispo PJM, Shanbhag SS, Gilmore MS, Chodosh J. The persistent dilemma of microbial keratitis: Global burden, diagnosis, and antimicrobial resistance. Surv Ophthalmol. 2019;64(3):255–71.

3. Tan SZ, Walkden A, Au L, Fullwood C, Hamilton A, Qamruddin A, Armstrong M, Brahma AK, Carley F. Twelve-year analysis of microbial keratitis trends at a UK tertiary hospital. Eye (Lond). 2017;31(8):1229–36.

4. Ting DSJ, Ho CS, Cairns J, Elsahn A, Al-Aqaba M, Boswell T, Said DG, Dua HS. 12-year analysis of incidence, microbiological profiles and in vitro antimicrobial susceptibility of infectious keratitis: the Nottingham Infectious Keratitis Study. Br J Ophthalmol. 2021;105(3):328–33.

5. Asbell PA, Sanfilippo CM, Sahm DF, DeCory HH. Trends in Antibiotic Resistance Among Ocular Microorganisms in the United States From 2009 to 2018. JAMA Ophthalmol. 2020;138(5):439–50.

6. Ting DSJ, Settle C, Morgan SJ, Baylis O, Ghosh S. A 10-year analysis of microbiological profiles of microbial keratitis: the North East England Study. Eye (Lond). 2018;32(8):1416–7.

7. Lee JW, Somerville T, Kaye SB, Romano V. Staphylococcus aureus Keratitis: Incidence, Pathophysiology, Risk Factors and Novel Strategies for Treatment. J Clin Med. 2021;10(4).

8. Ting DSJ, Henein C, Said DG, Dua HS. Photoactivated chromophore for infectious keratitis-corneal cross-linking (PACK-CXL): A systematic review and meta-analysis. Ocul Surf. 2019;17(4):624–34.

9. Tabatabaei SA, Soleimani M, Behrouz MJ, Torkashvand A, Anvari P, Yaseri M. A randomized clinical trial to evaluate the usefulness of amniotic membrane transplantation in bacterial keratitis healing. Ocul Surf. 2017;15(2):218–26.

10. Ting DSJ, Henein C, Said DG, Dua HS. Amniotic membrane transplantation for infectious keratitis: a systematic review and meta-analysis. Sci Rep. 2021;11(1):13007.

11. Khor WB, et al. The Asia Cornea Society Infectious Keratitis Study: A Prospective Multicenter Study of Infectious Keratitis in Asia. Am J Ophthalmol. 2018;195:161–70.

12. Ting DSJ, Bignardi G, Koerner R, Irion LD, Johnson E, Morgan SJ, Ghosh S. Polymicrobial Keratitis With Cryptococcus curvatus, Candida parapsilosis, and Stenotrophomonas maltophilia After Penetrating Keratoplasty: A Rare Case Report With Literature Review. Eye Contact Lens. 2019;45(2):e5–e10.

13. Khoo P, Cabrera-Aguas MP, Nguyen V, Lahra MM, Watson SL. Microbial keratitis in Sydney, Australia: risk factors, patient outcomes, and seasonal variation. Graefes Arch Clin Exp Ophthalmol. 2020;258(8):1745–55.

14. Asbell PA, Sanfilippo CM, Pillar CM, DeCory HH, Sahm DF, Morris TW. Antibiotic Resistance Among Ocular Pathogens in the United States: Five-Year Results From the Antibiotic Resistance Monitoring in Ocular Microorganisms (ARMOR) Surveillance Study. JAMA Ophthalmol. 2015;133(12):1445–54.

15. Nithya V, Rathinam S, Siva Ganesa Karthikeyan R, Lalitha P. A ten year study of prevalence, antimicrobial susceptibility pattern, and genotypic characterization of Methicillin resistant Staphylococcus aureus causing ocular infections in a tertiary eye care hospital in South India. Infect Genet Evol. 2019;69:203–10.

16. Kaye S, et al. Bacterial susceptibility to topical antimicrobials and clinical outcome in bacterial keratitis. Invest Ophthalmol Vis Sci. 2010;51(1):362–8.

17. Lalitha P, et al. Relationship of in vitro susceptibility to moxifloxacin and in vivo clinical outcome in bacterial keratitis. Clin Infect Dis. 2012;54(10):1381–7.

18. Ting DSJ, McKenna M, Sadiq SN, Martin J, Mudhar HS, Meeney A, Patel T. Arthrographis kalrae Keratitis Complicated by Endophthalmitis: A Case Report With Literature Review. Eye Contact Lens. 2020;46(6):e59–e65.

19. Ting DSJ, Cairns J, Gopal BP, Ho CS, Krstic L, Elsahn A, Lister M, Said DG, Dua HS. Risk Factors, Clinical Outcomes and Prognostic Factors of Bacterial Keratitis: The Nottingham Infectious Keratitis Study. Front Med (Lausanne). 2021; doi: 10.3389/fmed.2021.715118.

20. Hutchings MI, Truman AW, Wilkinson B. Antibiotics: past, present and future. Curr Opin Microbiol. 2019;51:72–80.

21. Hancock RE, Lehrer R. Cationic peptides: a new source of antibiotics. Trends Biotechnol. 1998;16(2):82–8.

22. Mookherjee N, Anderson MA, Haagsman HP, Davidson DJ. Antimicrobial host defence peptides: functions and clinical potential. Nat Rev Drug Discov. 2020.

23. Mohammed I, Said DG, Dua HS. Human antimicrobial peptides in ocular surface defense. Prog Retin Eye Res. 2017;61:1–22.

24. Kolar SS, McDermott AM. Role of host-defence peptides in eye diseases. Cell Mol Life Sci. 2011;68(13):2201–13.

25. Ting DSJ, Beuerman RW, Dua HS, Lakshminarayanan R, Mohammed I. Strategies in Translating the Therapeutic Potentials of Host Defense Peptides. Front Immunol. 2020;11:983.

26. Hancock RE, Sahl HG. Antimicrobial and host-defense peptides as new anti-infective therapeutic strategies. Nat Biotechnol. 2006;24(12):1551–7.

27. McIntosh RS, Cade JE, Al-Abed M, Shanmuganathan V, Gupta R, Bhan A, Tighe PJ, Dua HS. The spectrum of antimicrobial peptide expression at the ocular surface. Invest Ophthalmol Vis Sci. 2005;46(4):1379–85.

28. Haynes RJ, Tighe PJ, Dua HS. Innate defence of the eye by antimicrobial defensin peptides. Lancet. 1998;352(9126):451–2.

29. Haynes RJ, Tighe PJ, Dua HS. Antimicrobial defensin peptides of the human ocular surface. Br J Ophthalmol. 1999;83(6):737–41.

30. Mohammed I, Mohanty D, Said DG, Barik MR, Reddy MM, Alsaadi A, Das S, Dua HS, Mittal R. Antimicrobial peptides in human corneal tissue of patients with fungal keratitis. Br J Ophthalmol. 2020; doi: 10.1136/bjophthalmol-2020-316329.

31. Otri AM, Mohammed I, Abedin A, Cao Z, Hopkinson A, Panjwani N, Dua HS. Antimicrobial peptides expression by ocular surface cells in response to Acanthamoeba castellanii: an in vitro study. Br J Ophthalmol. 2010;94(11):1523–7.

32. Abedin A, Mohammed I, Hopkinson A, Dua HS. A novel antimicrobial peptide on the ocular surface shows decreased expression in inflammation and infection. Invest Ophthalmol Vis Sci. 2008;49(1):28–33.

33. Otri AM, Mohammed I, Al-Aqaba MA, Fares U, Peng C, Hopkinson A, Dua HS. Variable expression of human Beta defensins 3 and 9 at the human ocular surface in infectious keratitis. Invest Ophthalmol Vis Sci. 2012;53(2):757–61.

34. Huang LC, Jean D, Proske RJ, Reins RY, McDermott AM. Ocular surface expression and in vitro activity of antimicrobial peptides. Curr Eye Res. 2007;32(7-8):595–609.

35. Gordon YJ, Huang LC, Romanowski EG, Yates KA, Proske RJ, McDermott AM. Human cathelicidin (LL-37), a multifunctional peptide, is expressed by ocular surface epithelia and has potent antibacterial and antiviral activity. Curr Eye Res. 2005;30(5):385–94.

36. Li J, Koh JJ, Liu S, Lakshminarayanan R, Verma CS, Beuerman RW. Membrane Active Antimicrobial Peptides: Translating Mechanistic Insights to Design. Front Neurosci. 2017;11:73.

37. Fjell CD, Hiss JA, Hancock RE, Schneider G. Designing antimicrobial peptides: form follows function. Nat Rev Drug Discov. 2011;11(1):37–51.

38. Boman HG, Wade D, Boman IA, Wahlin B, Merrifield RB. Antibacterial and antimalarial properties of peptides that are cecropin-melittin hybrids. FEBS Lett. 1989;259(1):103–6.

39. Wei XB, Wu RJ, Si DY, Liao XD, Zhang LL, Zhang RJ. Novel Hybrid Peptide Cecropin A (1-8)-LL37 (17-30) with Potential Antibacterial Activity. Int J Mol Sci. 2016;17(7).

40. Willcox MD, Hume EB, Aliwarga Y, Kumar N, Cole N. A novel cationic-peptide coating for the prevention of microbial colonization on contact lenses. J Appl Microbiol. 2008;105(6):1817–25.

41. Willcox MD, Chen R, Kalaiselvan P, Yasir M, Rasul R, Kumar N, Dutta D. The development of an antimicrobial contact lens - From the laboratory to the clinic. Curr Protein Pept Sci. 2020;21(4):357–68.

42. Mohammed I, Said DG, Nubile M, Mastropasqua L, Dua HS. Cathelicidin-Derived Synthetic Peptide Improves Therapeutic Potential of Vancomycin Against Pseudomonas aeruginosa. Front Microbiol. 2019;10:2190.

43. Kampshoff F, Willcox MDP, Dutta D. A Pilot Study of the Synergy between Two Antimicrobial Peptides and Two Common Antibiotics. Antibiotics (Basel). 2019;8(2).

44. Saugar JM, Rodriguez-Hernandez MJ, de la Torre BG, Pachon-Ibanez ME, Fernandez-Reyes M, Andreu D, Pachon J, Rivas L. Activity of cecropin A-melittin hybrid peptides against colistin-resistant clinical strains of Acinetobacter baumannii: molecular basis for the differential mechanisms of action. Antimicrob Agents Chemother. 2006;50(4):1251–6.

45. McDermott AM. The role of antimicrobial peptides at the ocular surface. Ophthalmic Res. 2009;41(2):60–75.

46. Wang G. Structures of human host defense cathelicidin LL-37 and its smallest antimicrobial peptide KR-12 in lipid micelles. J Biol Chem. 2008;283(47):32637–43.

47. Krishnakumari V, Nagaraj R. Binding of peptides corresponding to the carboxy-terminal region of human-β-defensins-1-3 with model membranes investigated by isothermal titration calorimetry. Biochim Biophys Acta. 2012;1818(5):1386–94.

48. Dhople V, Krukemeyer A, Ramamoorthy A. The human beta-defensin-3, an antibacterial peptide with multiple biological functions. Biochim Biophys Acta. 2006;1758(9):1499–512.

49. Strahl H, Errington J. Bacterial Membranes: Structure, Domains, and Function. Annu Rev Microbiol. 2017;71:519–38.

50. Mahlapuu M, Hakansson J, Ringstad L, Bjorn C. Antimicrobial Peptides: An Emerging Category of Therapeutic Agents. Front Cell Infect Microbiol. 2016;6:194.

51. Kolar SSN, Luca V, Baidouri H, Mannino G, McDermott AM, Mangoni ML. Esculentin-1a(1-21)NH2: a frog skin-derived peptide for microbial keratitis. Cell Mol Life Sci. 2015;72(3):617–27.

52. Clemens LE, Jaynes J, Lim E, Kolar SS, Reins RY, Baidouri H, Hanlon S, McDermott AM, Woodburn KW. Designed Host Defense Peptides for the Treatment of Bacterial Keratitis. Invest Ophthalmol Vis Sci. 2017;58(14):6273–81.

53. Dutta D, Zhao T, Cheah KB, Holmlund L, Willcox MDP. Activity of a melimine derived peptide Mel4 against Stenotrophomonas, Delftia, Elizabethkingia, Burkholderia and biocompatibility as a contact lens coating. Cont Lens Anterior Eye. 2017;40(3):175–83.

54. Mayandi V, et al. Rational Substitution of ɛ-Lysine for α-Lysine Enhances the Cell and Membrane Selectivity of Pore-Forming Melittin. J Med Chem. 2020;63(7):3522–37.

55. Kapoor G, Saigal S, Elongavan A. Action and resistance mechanisms of antibiotics: A guide for clinicians. J Anaesthesiol Clin Pharmacol. 2017;33(3):300–5.

56. Farkouh A, Frigo P, Czejka M. Systemic side effects of eye drops: a pharmacokinetic perspective. Clin Ophthalmol. 2016;10:2433–41.

57. Seo J, et al. Multiscale reverse engineering of the human ocular surface. Nat Med. 2019;25(8):1310–8.

58. Girgis DO, Sloop GD, Reed JM, O’Callaghan RJ. A new topical model of Staphylococcus corneal infection in the mouse. Invest Ophthalmol Vis Sci. 2003;44(4):1591–7.

59. Lakshminarayanan R, et al. Branched peptide, B2088, disrupts the supramolecular organization of lipopolysaccharides and sensitizes the Gram-negative bacteria. Sci Rep. 2016;6:25905.

60. Mishra B, Reiling S, Zarena D, Wang G. Host defense antimicrobial peptides as antibiotics: design and application strategies. Curr Opin Chem Biol. 2017;38:87–96.

61. Smith JE. Erythrocyte membrane: structure, function, and pathophysiology. Vet Pathol. 1987;24(6):471–6.

62. Tavassoli S, Nayar G, Darcy K, Grzeda M, Luck J, Williams OM, Tole D. An 11-year analysis of microbial keratitis in the South West of England using brain-heart infusion broth. Eye (Lond). 2019;33(10):1619–25.

63. Clinical & Laboratory Standards Institute (CLSI). Methods for dilution antimicrobial susceptibility tests for bactera that grow aerobically, 11th edition. 2019.

64. Lin S, et al. Symmetrically Substituted Xanthone Amphiphiles Combat Gram-Positive Bacterial Resistance with Enhanced Membrane Selectivity. J Med Chem. 2017;60(4):1362–78.

65. Kilkenny C, Browne WJ, Cuthill IC, Emerson M, Altman DG. Improving bioscience research reporting: the ARRIVE guidelines for reporting animal research. PLoS Biol. 2010;8(6):e1000412.

66. Aung TT, Yam JK, Lin S, Salleh SM, Givskov M, Liu S, Lwin NC, Yang L, Beuerman RW. Biofilms of Pathogenic Nontuberculous Mycobacteria Targeted by New Therapeutic Approaches. Antimicrob Agents Chemother. 2016;60(1):24–35.

67. Schneider CA, Rasband WS, Eliceiri KW. NIH Image to ImageJ: 25 years of image analysis. Nature Methods 2012;9(7):671–5.

