## Supplementary material for "Hybrid Derivative of Cathelicidin and Human Beta Defensin-2 Against Gram-Positive Bacteria: A Novel Approach for the Treatment of Bacterial Keratitis": Table 2

**Table 2.** A summary of the minimum inhibitory concentrations (MICs) of various antibiotics and synthetic human-derived hybrid host defense peptides. All experiments were performed in full-strength cationic Muller-Hinton broth (i.e. MHB-2) and in the absence or presence of physiological tear salt concentration (150mM NaCl). The MIC values are presented in μg/ml (and μM in bracket). When the MIC level was ≥200 μg/ml in the absence of salt, the peptide was not subjected to testing in the presence of salt.

| **Class of agent** | **Agents**  μg/ml (μM) | ***S. aureus***  **SH1000** | | | ***S. aureus* ATCC 29213** | | **MRSA-OS** | | **MRSA**  **ATCC 43300** | | ***S. epidermidis***  **ATCC 12228** | | ***P. aeruginosa***  **PAO1-L** | | | ***P. aeruginosa***  **PA-OS** | | |
| --- | --- | --- | --- | --- | --- | --- | --- | --- | --- | --- | --- | --- | --- | --- | --- | --- | --- | --- |
|  |  | *0mM* | *150mM* | *0mM* | | *150mM* | *0mM* | *150mM* | *0mM* | *150mM* | *0mM* | *150mM* | | *0mM* | *150mM* | | *0mM* | *150mM* |
| Antibiotics | Amikacin | 1.25  (2.13) | 2.5  (4.27) | - | | - | 10  (17.1) | 10  (17.1) | - | - | - | - | | 0.63  (1.08) | 1.25  (2.13) | | 0.63  (1.08) | 1.25  (2.13) |
|  | Levofloxacin | 0.31  (0.86) | 0.31  (0.86) | - | | - | 0.31  (0.86) | 0.31  (0.86) | - | - | - | - | | 0.31  (0.86) | 0.31  (0.86) | | 0.31  (0.86) | 0.31  (0.86) |
| Full-length peptide | LL37 | 25  (5.6) | - | - | | - | 25  (5.6) | - | - | - | - | - | | 50  (11.1) | - | | - | - |
|  | HBD2 | 100  (23.1) | - | - | | - | 100  (23.1) | - | - | - | - | - | | 100  (23.1) | - | | - | - |
|  | HBD3 | 100  (19.4) | - | - | | - | 100  (19.4) | - | - | - | - | - | | 100  (19.4) | - | | - | - |
| Single linear HDPs | Ca12 | >200  (>127.2) | - | - | | - | >200  (>127.2) | - | - | - | - | - | | 200  (127.2) | - | | - | - |
|  | BD2-6 | >200  (>283.3) | - | - | | - | >200  (>283.3) | - | - | - | - | - | | >200  (>283.3) | - | | - | - |
|  | BD3-10 | >200  (>154.9) |  |  | |  | >200  (>154.9) |  |  |  |  |  | | >200  (>154.9) |  | | - | - |
| First-generation hybrid HDPs | DD12 | >200  (>115) | - | - | | - | >200  (>115) | - | - | - | - | - | | >200  (>115) | - | | - | - |
|  | DD13 | >200  (>92.3) | - | - | | - | >200  (>92.3) | - | - | - | - | - | | >200  (>92.3) | - | | - | - |
|  | DD32 | >200  (>101) | - | - | | - | >200  (>101) | - | - | - | - | - | | >200  (>101) | - | | - | - |
|  | CaD1 | >200  (>94.7) | - | - | | - | >200  (>94.7) | - | - | - | - | - | | >200 (>94.7) | - | | - | - |
|  | CaD2 | >200  (>88.5) | - | - | | - | >200  (>88.5) | - | - | - | - | - | | >200  (>88.5) | - | | - | - |
|  | CaD3 | >200  (>91.8) | - | - | | - | >200  (>91.8) | - | - | - | - | - | | >200  (>91.8) | - | | - | - |
| Second-generation hybrid HDPs | CaD21 | 200  (89.8) | - | - | | - | >200  (>89.8) | - | - | - | - | - | | 100  (44.9) | 100  (44.9) | | 200  (89.8) | - |
|  | CaD22 | 100  (43.5) | >200  (>87.0) | - | | - | >200  (>87.0) | - | - | - | - | - | | 200  (87.0) | - | | 200  (87.0) | - |
|  | CaD23 | 12.5  (5.2) | 25  (10.4) | 25  (10.4) | | 50  (20.8) | 25  (10.4) | 100  (41.7) | 25  (10.4) | 100  (41.7) | 12.5  (5.2) | 12.5  (5.2) | | 50  (20.8) | 50  (20.8) | | 25  (10.4) | 50  (20.8) |

ATCC = American Type Culture Collection; MRSA-OS = Methicillin-resistant *S. aureus*; OS = Ocular surface

MIC refers to the lowest concentration of antibiotic / peptide that prevents any visible bacterial growth after 24 hours of incubation with treatment. Data represent the mean of two biological duplicate from two to three independent experiments.
