## Supplementary material for "Hybrid Derivative of Cathelicidin and Human Beta Defensin-2 Against Gram-Positive Bacteria: A Novel Approach for the Treatment of Bacterial Keratitis": Table 3

**Table 3.** Summary of the cytotoxicity, cell viability and hemolytic results of antibiotics and synthetic peptides (in µg/ml concentration). The cytotoxicity and cell viability results were obtained after 3 hours of treatment whereas hemolytic effect of treatment was examined after 1 hour of treatment.

| **Types** | **Agents** | **LC_50_** | **L_max_ (%)** | **IC_50_** | **I_max_ (%)** | **HC_50_** | **H_max_ (%)** |
| --- | --- | --- | --- | --- | --- | --- | --- |
| Antibiotics | Amikacin | >200 | 0.0 (0.1) | >200 | 100.1 (7.3) | >200 | 0.0 |
|  | Levofloxacin | >200 | 1.2 (0.7) | >200 | 108.0 (2.8) | >200 | 0 (0.0) |
| First-generation peptides | DD12 | - | - | - | - | - | - |
|  | DD13 | - | - | - | - | - | - |
|  | DD32 | - | - | - | - | - | - |
|  | CaD1 | - | - | - | - | - | - |
|  | CaD2 | >200 | 2.9 (2.2) | >200 | 1.9 (7.3) | - | - |
|  | CaD3 | - | - | - | - | - | - |
| Second-generation peptides | CaD21 | - | - | >200 | 43.9 (1.0) | - | - |
|  | CaD22 | - | - | >200 | 34.3 (8.2) | - | - |
|  | CaD23 | >200 | 26.6 (6.5) | 54.6 (11.7) | 69.6 (7.8) | >200 | 7.1 (3.0) |

LC_50_ = Concentration of treatment causing 50% cytotoxicity; L_max_ (%) = Percentage of cytotoxicity at 200 µg/ml treatment concentration; IC_50_ = Concentration of treatment causing 50% inhibition of cell viability; I_max_ (%) = Percentage of inhibition of cell viability at 200 µg/ml treatment concentration;

HC_50_ = Concentration of treatment causing 50% hemolysis; H_max_ (%) = Percentage of hemolysis at 200 µg/ml treatment concentration

Results are presented in mean (SD) of two independent experiments performed in biological duplicate. Toxicity results of some peptides were missing because their antimicrobial efficacy was poor and hence toxicity was not determined.
