## Supplementary material for "Hybrid Derivative of Cathelicidin and Human Beta Defensin-2 Against Gram-Positive Bacteria: A Novel Approach for the Treatment of Bacterial Keratitis": Table 4

**Table 4.** Summary of therapeutic index of CaD23 determined based on cytotoxicity, cell viability, and hemolytic results. The minimum inhibitory concentration (MIC) values, based on the results in the presence of full-strength cationic Muller-Hinton broth (caMHB) without salt, and therapeutic index (TI) values are presented in mean values.

| **μg/ml** | ***S. aureus***  **SH1000** | | ***S. aureus***  **ATCC29213** | | **MRSA-OS** | | **MRSA**  **ATCC43300** | | ***S. epidermidis***  **ATCC12228** | | ***P. aeruginosa* PA01-L** | | ***P. aeruginosa***  **PA-OS** | |
| --- | --- | --- | --- | --- | --- | --- | --- | --- | --- | --- | --- | --- | --- | --- |
|  | **MIC** | **TI** | **MIC** | **TI** | **MIC** | **TI** | **MIC** | **TI** | **MIC** | **TI** | **MIC** | **TI** | **MIC** | **TI** |
| Based on LC_50_ (= >200) | 12.5 | >16 | 25 | >8 | 25 | >8 | 50 | >4 | 12.5 | >16 | 50 | >4 | 25 | >8 |
| Based on IC_50_  (= 55) | 12.5 | 4.4 | 25 | 2.2 | 25 | 2.2 | 50 | 1.1 | 12.5 | 4.4 | 50 | 1.1 | 25 | 2.2 |
| Based on HC_50_ (= >200) | 12.5 | >16 | 25 | >8 | 25 | >8 | 50 | >4 | 12.5 | >16 | 50 | >4 | 25 | >8 |

MRSA = Methicillin-resistant *Staphylococcus aureus*; OS = Ocular surface (clinical isolate)

LC_50_ = Concentration of treatment causing 50% cytotoxicity; IC_50_ = Concentration of treatment causing 50% inhibition of cell viability; HC_50_ = Concentration of treatment causing 50% hemolysis
